# Comparative Analysis of Whole Transcriptome Single-Cell Sequencing Technologies in Complex Tissues

**DOI:** 10.1101/2023.07.03.547464

**Authors:** Stefan Salcher, Isabel Heidegger, Gerold Untergasser, Georgios Fotakis, Alexandra Scheiber, Agnieszka Martowicz, Asma Noureen, Anne Krogsdam, Christoph Schatz, Georg Schäfer, Zlatko Trajanoski, Dominik Wolf, Sieghart Sopper, Andreas Pircher

## Abstract

The development of single-cell omics tools has enabled scientists to study the tumor microenvironment (TME) in unprecedented detail. However, each of the different techniques may have its unique strengths and limitations. Here we directly compared two commercially available high-throughput single-cell RNA sequencing (scRNA-seq) technologies - droplet-based 10X Chromium *vs.* microwell-based BD Rhapsody - using paired samples from patients with localized prostate cancer (PCa) undergoing a radical prostatectomy.

Although high technical consistency was observed in unraveling the whole transcriptome, the relative abundance of cell populations differed. Cells with low-mRNA content such as T cells were underrepresented in the droplet-based system, at least partly due to lower RNA capture rates. In contrast, microwell based scRNA-seq recovered less cells of epithelial origin. Moreover, we discovered platform-dependent variabilities in mRNA quantification and cell-type marker annotation. Overall, our study provides important information for selection of the appropriate scRNA-seq platform and for the interpretation of published results.

**SYNOPSIS:**

- Comparison of scRNA-seq protocols uncovers disparities in RNA-to-library conversion
- Microwell-based scRNA-seq technology excels in capturing low-mRNA content cells
- Biased transcriptomes due to gene specific RNA detection efficacies by both platforms
- The study guides in informed scRNA-seq platform selection and data interpretation

## INTRODUCTION

In the past decade a variety of high-dimensional single-cell omics tools have been developed and optimized at exponential pace, providing unprecedented opportunities to deconvolute tissue composition in various research fields including the tumor microenvironment (TME). This enabled the discovery of novel cell types with previously unknown functions across different diseases including various malignancies (Azizi *et al*, 2018; Goveia *et al*, 2020a; Guo *et al*, 2018; Lambrechts *et al*, 2018; Li *et al*, 2019; Zhang *et al*, 2019a). In contrast to other single-cell technologies such as fluorescence based or mass spectral flow cytometry, this enabled the unbiased discovery of novel cell types and distinct functional states across multicellular tissues. Since the first description in 2009 (Tang *et al*, 2009), several scRNA-seq platforms have been introduced to refine scRNA-seq throughput, sensitivity, precision, and costs. These advances paved the way to broadly apply the technology to map cellular and molecular complexity of the TME and to characterize diverse cell types in cancer (Azizi *et al*., 2018; Goveia *et al*, 2020b; Li *et al*., 2019; Zheng *et al*, 2017a). ScRNA-seq is used to quantify the individual cell-specific transcriptome at single-cell level. Gene expression levels are denoted by the sequenced reads and the unique transcriptomes individual cell types express, which are then presented as a data matrix. Hence, a typical scRNA-seq workflow would encompass single-cell isolation and capture, sample barcoding, mRNA reverse transcription, cDNA amplification and cDNA library preparation, next-generation sequencing (NGS), and computational data processing.

Different scRNA-seq strategies have been developed to generate either full-length cDNA (for full-length sequencing) or cDNA with a unique molecular identifier (UMI) at the 3′ or 5′ end (for 3′/5′ sequencing). Full-length sequencing protocols like SMART-seq (Ramskold *et al*, 2020), as well as the improved version SMART-seq2 (Picelli *et al*, 2013), generate reads across the entire length of genes by template switching-based PCR. The high sensitivity to detect gene expression enables various downstream analyses of specific cell types, tissue composition, spliced transcript variants, isoforms, or allele-specific gene expression patterns (See *et al*, 2019). Full-length sequencing is low-throughput, relatively expensive, and struggles with a gene length bias akin to bulk RNA-seq data. Genes with shorter lengths generally exhibit lower read counts and a higher drop-out rate (Phipson *et al*, 2017). “Molecular tagging” with UMIs (Kivioja *et al*, 2011) used in 3′/5′ sequencing mitigated biases from downstream PCR amplification and enables digital counting of absolute numbers of each transcript per individual cell. In addition, UMIs allow sample multiplexing to improve gene expression quantification and throughput, and thus reduce overall costs (See *et al*., 2019). Although the sensitivity is inferior to full-length sequencing, tag-based sequencing methods are more suitable for quantification purposes (e.g. cell typing), this is particularly true when considering the simultaneous sampling of tens of thousands of cells (Hedlund & Deng, 2018; Kalisky *et al*, 2018; See *et al*., 2019).

The technology used to capture a single cell for RNA sequencing determines the number of isolated cells, whether there is a biased or unbiased selection of cells, the integrity and purity of the cells, and lastly the costs of the experiment. For biological samples containing only few cells, manual low throughput methods, such as laser capture microdissection (LCM), limiting dilution of cell suspensions, or manual cell picking with micromanipulators are applicable. To increase throughput, FACS is suitable to analyze, sort, and enrich single cells. Still, cell numbers remain limited and downstream library preparation is laborious. Thus, distinctive small-volume microfluidic droplet-based 3′/5′ sequencing technologies are primarily applied as an ultra-high-throughput, unbiased solution (Zhang *et al*, 2019b). Currently, the prevalent droplet-based scRNA-seq systems are inDrop (Klein *et al*, 2015), Drop-seq (Macosko *et al*, 2015), and the commercially available platform 10X Chromium (10X Genomics, USA) (Zheng *et al*, 2017b). Each of these three technologies generates microfluidic droplets to capture single cells together utilizing on-bead primers with unique barcodes. Besides the droplet-based methods, Seqwell (Gierahn *et al*, 2017), Microwell-seq (Han *et al*, 2018), Cyto-seq (Fan *et al*, 2015), as well as the commercialized platforms Singleron GEXSCOPE (Singleron Biotechnologies, China) and BD Rhapsody (Becton Dickinson, USA) (Shum *et al*, 2019) apply microwell arrays into which individual cells are loaded together with barcoded magnetic beads. Upon cell lysis, the mRNA content of each cell is captured, reverse transcribed into cDNA, and amplified for sequencing-library generation through oligo-dT priming procedures.

These tools are helping scientists to sequence and study RNA in unprecedented detail, but each technique has its own strengths and inherent limitations as described in several comparative studies (Chen *et al*, 2021b; Colino-Sanguino *et al*, 2023; Gao *et al*, 2020; Mereu *et al*, 2020; Natarajan *et al*, 2019; Wang *et al*, 2021; Yamawaki *et al*, 2021; Zhang *et al*., 2019b; Ziegenhain *et al*, 2017). In addition, our recent scRNA-seq study on lung cancer, in which we integrated scRNA-seq data from 19 studies and 21 datasets, comprising 505 samples from 298 patients (Salcher *et al,* 2022) revealed that low-mRNA content cells are frequently dismissed due to technical issues. The integrated datasets were generated using various scRNA-seq platforms, including 10X Chromium and BD Rhapsody. Lung cancer tissue resident neutrophils, characterized by exceptionally low mRNA content, were predominantly detected in the dataset generated with BD Rhapsody, while they were underrepresented or even absent in datasets generated with 10X Chromium or other platforms (Salcher *et al.,* 2022). Using this substantial advantage of microwell-based scRNA-seq, we recently also characterized previously unrecognized tissue resident low-mRNA content neutrophils in human livers (Hautz *et al*, 2023).

These performance differences have been described using either rather artificial cell systems (Mereu *et al*., 2020) or comparisons of data from different studies (Salcher *et al*., 2022) but a rigorous side by side comparison of primary samples covering the complete cellular heterogeneity within the tissue has not been performed. Thus, we performed a systematic comparison of two commercially available high-throughput scRNA-seq technology platforms, i.e. the droplet-based 10X Chromium and the microwell-based BD Rhapsody platform, using paired surgically resected prostate cancer (PCa) and corresponding benign tissues, in order to derive the necessary information for the planning of scRNA-seq experiments and the interpretation of the results.

## RESULTS

### Droplet-vs. microwell-based single-cell RNA sequencing of PCa and benign prostate tissues

To directly compare the capability to deconvolute the TME and benign tissue composition at single-cell resolution of the most abundant cancer in men, we dissociated six freshy isolated tissue samples from treatment naïve localized PCa patients undergoing a radical prostatectomy (3 tumor tissues samples and 3 corresponding benign prostate tissues samples) and subjected the obtained single-cell suspensions to 3’WTA scRNA-seq analysis using the 10X Chromium as well as the BD Rhapsody platforms, respectively (the experimental setup is illustrated in Figure 1A; selection criteria of tissue samples obtained from fresh radical prostatectomy (RP) specimens to perform scRNA-seq analysis have been described previously (Heidegger *et al*, 2022); the detailed scRNAseq workflow of both platforms is shown in Figure S1).

**Figure 1.**
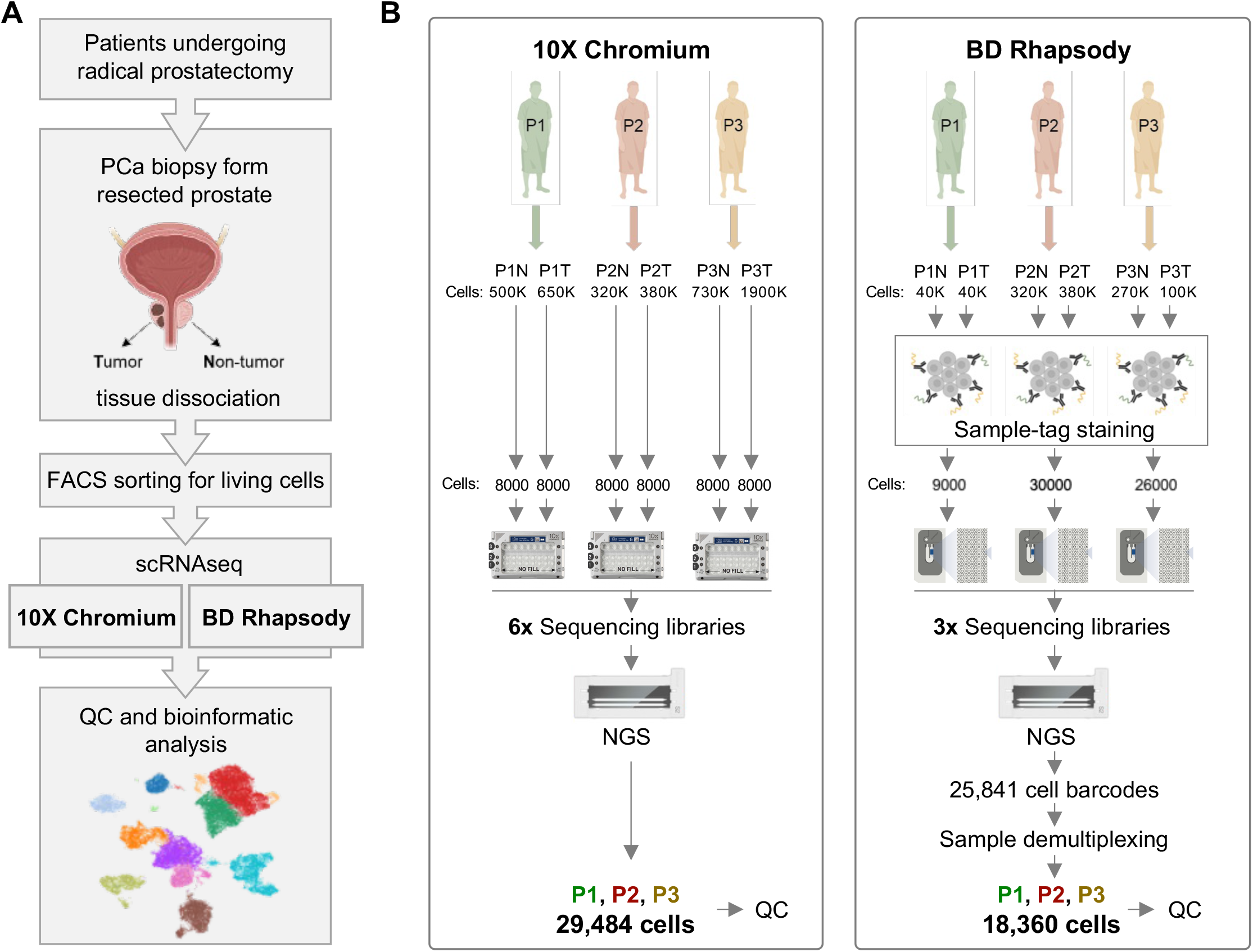
Experimental setup. (A) Summary of the analysis workflow. (B) scRNA-seq datasets were generated from freshly isolated benign prostate (n=3) and PCa (n=3) tissues using the 10X Chromium and BD Rhapsody platforms, respectively. Cell numbers are shown for each step starting with number of isolated cells from each sample used for the two platforms. In the 10X Chromium dataset 29,484 cells (cells with >100 genes expressed in ≥3 cells) were detected and subjected to quality control processing. In the BD Rhapsody dataset 25,841 cell barcodes were identified and 21,196 cell barcodes with sample-tag information could be recovered during sample demultiplexing. Thereof, 18,360 cells (cells with >100 genes expressed in ≥3 cells) were subjected to quality control processing. See also **Figure S1**.

To investigate the capability of the 10X Chromium protocol, we generated and sequenced six WTA index libraries from a total of 48,000 cells, with 8,000 cells per sample. After pre-processing the raw sequencing data using the 10X Chromium Cell Ranger software, we detected 29,484 cells (cells with >100 genes expressed in ≥3 cells), corresponding to a capture rate of ∼60% (including multiplets; Figure 1B).

In the BD Rhapsody workflow, we used the BD Rhapsody Single Cell Multiplexing kit (SMK) to process multiple samples simultaneously. Following the process of labeling single cells derived from both benign and tumor tissue using sample-tag antibody staining, we generated a separate sequencing library for each of the three individual patients. In contrast to 10X Chromium, the microwell-based BD Rhapsody workflow enables the quantification of single living cells captured together with a bead within the system. Hence, this detailed information about the quality and number of captured single cells, as well as the number of captured multiplets, enables a more precise calculation of the desired sequencing depth per cell, especially when multiple samples are combined in one sequencing run. Thus, it is possible to use a predefined number of cells for WTA index library preparation and to store remaining beads for later use. The capability of the BD Rhapsody platform for sample-multiplexing and the flexibility to adjust cell numbers significantly contribute to cost reduction in library preparation and optimization of subsequent sequencing expenses. From the 65,000 deployed single-cells, 32,000 living cells including 5.5% multiplets could be captured together with a bead. Sequencing resulted in 25,841 detected cell barcodes. Still, just 21,196 cell barcodes with sample-tag information could be recovered during sample demultiplexing after pre-processing of the raw sequencing data using the BD Rhapsody WTA Analysis Pipeline app in the cloud-based Seven Bridges Genomics environment. Approximately 10-15% of “undetermined” cell barcodes with no sample-tag information had to be excluded from downstream analyses. Overall, we could depict 18,360 cells (cells with >100 genes expressed in ≥3 cells), resulting in an effective cell capture rate of ∼30% (excluding multiplets, Figure 1B).

In summary, 29,484 cells (10,557 benign (∼36%) and 18,927 tumor (∼64%)) acquired with 10X Chromium as well as 18,360 cells (7,145 benign (∼39%) and 11,215 tumor (∼61%)) obtained with BD Rhapsody (detected cells in each individual sample are detailed in Table S1), were subjected to quality control (QC) processing, a critical step during scRNA-seq data processing (Hwang *et al*, 2018).

### Microwell-based single-cell RNA sequencing results in elevated levels of mitochondrial transcripts

During QC user-defined thresholds for different metrics computed for each individual cell are applied to filter for “high-quality” cells to yield biologically meaningful results in subsequent down-stream analyses (Luecken & Theis, 2019). The three commonly used QC thresholds are the number of different genes detected in each cell, the total transcript count per cell (also known as the library size), as well as the ratio of reads mapped to mitochondrial DNA-encoded genes to the total number of reads mapped. These metrics have to be adjusted individually depending on the analyzed cell or tissue type, respectively (Ji & Sadreyev, 2019).

Here, we filtered for cells with > 200 and < 8000 detected genes (nFeatures), total transcript counts > 2000 (nCounts), and < 30% of mitochondrial transcripts (%MT) (Figure 2A). 16,111 cells sequenced with 10X Chromium and 10,155 cells sequenced with BD Rhapsody passed the quality control step (26,266 cells in total; Table S1). Although the overall cell recovery rates (cells passing QC) of the processed PCa and benign prostate samples were well comparable between both scRNA-seq platforms (BD Rhapsody: 55.3%, 10X Chromium: 54.6%; metrics for each individual sample are detailed in Table S1), we found substantial differences in individual QC metrics between the two platforms (Figure 2B). In all six individual samples the microwell-based BD Rhapsody protocol resulted in significantly less genes (nFeatures), a higher number of transcript counts per cell (nCounts), as well as a higher proportion of mitochondrial transcripts (%MT; Figure 2C, Figure S2).

**Figure 2.**
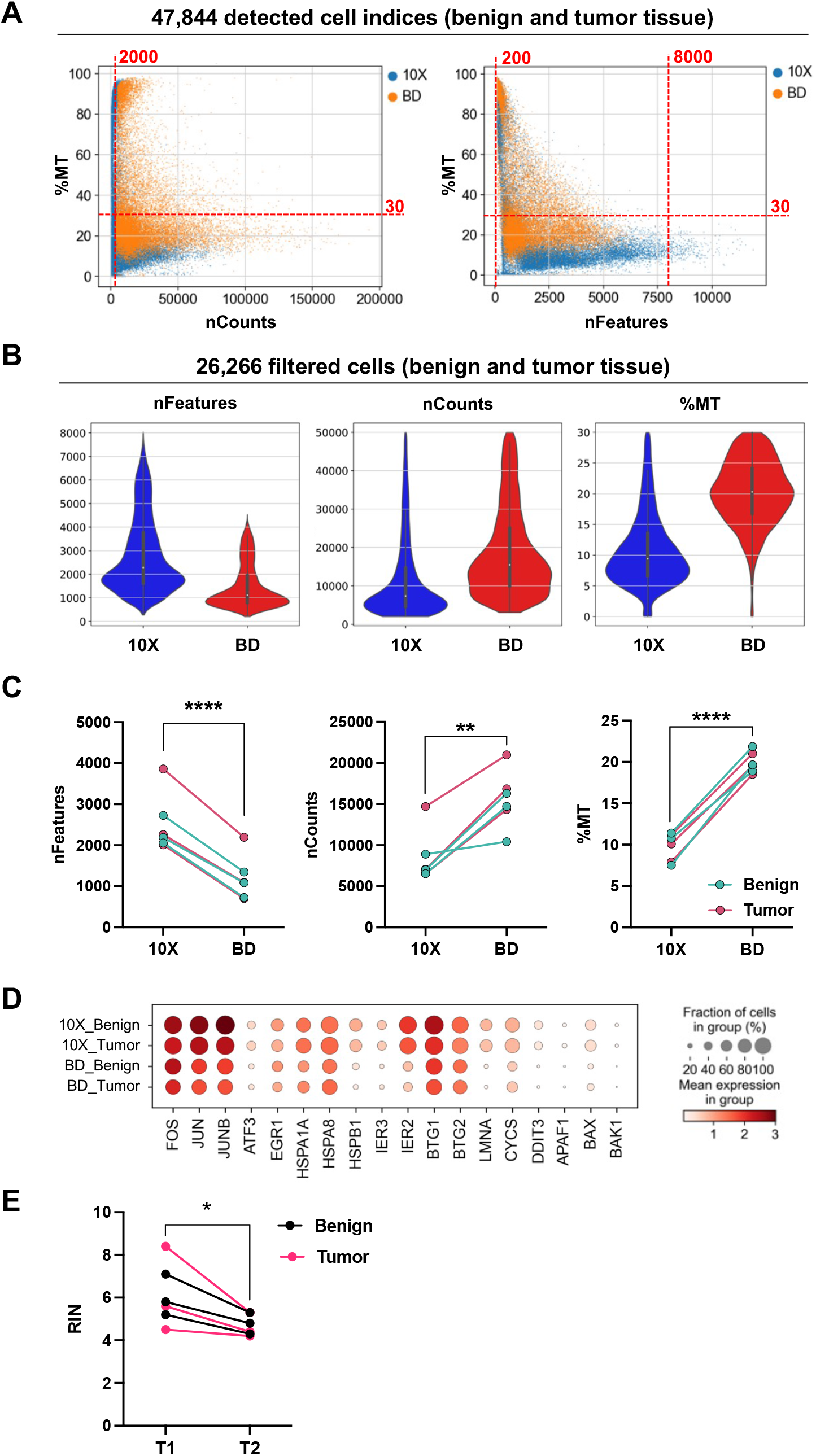
QC metrics in datasets generated with 10X Chromium and BD Rhapsody. (A) Correlation of %MT with nCounts and nFeatures quality metrics in data generated with 10X Chromium and BD Rhapsody using cells expressing > 100 genes (features). Applied cut-off values to filter for high quality cells are indicated (nCounts > 2000, nFeatures > 200 and < 8000, %MT < 30%). (B) nFeature, nCount, and %MT quality metrics in filtered cells derived from benign and PCa tumor tissues. (C) nFeature, nCount, and %MT levels in individual samples processed with 10X Chromium and BD Rhapsody (n=6; benign n=3, tumor n=3). Paired t-test, **p ≤ 0.01, ****p ≤ 0.0001. (D) Expression of stress-related transcripts in 10X Chromium and BD Rhapsody data generated from benign prostate and PCa samples. (E) RNA quality (RIN) before (T1) and after (T2) the sample-tag staining procedure in freshly isolated lung cancer (NSCLC) tumor tissues (n=6). Paired t-test, *p ≤ 0.05. See also **Figure S2** and **Figure S3**.

The fraction of reads mapped to mitochondrial DNA-encoded genes (%MT) represents a general indicator for cell stress, and thus a high level of mitochondrial transcripts has been associated with stressed, apoptotic, low-quality cells (Ilicic *et al,* 2016; Osorio & Ca*i, 2021*; *Zhao et al,* 2002). Although the proportion of mitochondrial counts per cell was significantly higher in the dataset generated with BD Rhapsody (mean %MT: 20.2) compared to 10X Chromium (mean %MT: 10.8), the expression of diverse stress-related transcripts did not markedly differ (Figure 2D). Cell stress-related transcripts, such as *IRE2*, *HSPB1*, *LMNA*, and *BAX*, were even expressed to a lesser extent in the BD Rhapsody dataset, in both benign prostate and PCa tissues, compared to 10X Chromium, respectively (Figure 2D).

### Sample-multiplexing by sample-tag antibody staining impairs RNA quality

In contrast to the droplet-based protocol, where each sample is processed separately, we utilized sample-tag antibodies in the BD Rhapsody workflow to enable multiplexing of multiple samples, including benign and tumor samples. A graphical representation of the detailed workflow from cell preparation to library construction for both platforms is outlined in Figure S1.

Next, we investigated whether the overall lengthier protocol of the microwell-based platform including the additional sample-tagging procedure may affect RNA quality in freshly isolated cell suspensions. For RNA quality benchmarking we used an additional cohort of freshly resected human lung tumors (n=3) as well as benign lung tissues (n=3). Following tissue dissociation, we subjected the obtained single-cell suspensions to the sample-tag staining procedure (20 min RT, 3x washing by 5 min centrifugation at 400 rpm). The BD Rhapsody sample-tag protocol significantly impaired RNA quality (RNA Integrity Number (RIN)) in multiple single-cell suspensions (n=6; Figure 2E). Importantly, transcripts associated with RNA decay did not exhibit elevated expression levels in BD Rhapsody data (Figure S3A), indicating that the RNA-degrading machinery is not actively induced during the scRNA-seq procedure. However, as the quality of RNA is a critical determinant of the reliability and accuracy of scRNA-seq data, any factors that compromise RNA quality may lead to biased results. Overall, these findings underscore the critical need for thorough evaluation and optimization of scRNA-seq protocols to ensure the reliability and validity of experimental results.

### The microwell-based platform captures significantly more mRNA molecules per cell

The number of genes detected in each cell (nFeatures) is linked to the applied sequencing depth (mean reads per cell per gene) - a measure of the available sequencing capacity spent for a single sample. The three sequencing libraries generated with BD Rhapsody were sequenced on the NovaSeq S1 flowcell (1.6 billion single reads; Illumuna) resulting in a mean coverage of ∼45,000 reads/cell. In contrast, the sequencing libraries prepared with the 10X Chromium system were sequenced on the NovaSeq S2 flowcell (4.1 billion single reads; Illumuna) resulting in a 1.6-fold higher mean coverage of ∼72,000 reads/cell (samples obtained from four PCa patients (∼64,000 cells) (Heidegger *et al*., 2022); three out of these four patients (∼48,000 cells) were processed in parallel with the BD Rhapsody platform). Consequently, the number of individual genes detected per cell was higher in the dataset obtained with 10X Chromium (Median nFeatures: BD Rhapsody ∼1,100, 10X Chromium ∼2,300, Figure 2B and Table S1). The elevated number of transcripts detected per cell was consistent over individual samples (Figure 2C, Figure S2A and Table S1) but did not markedly affect the number of quality cells after filtering (nFeatures > 200 and < 8000; Figure 2A and Figure S2B).

Remarkably, despite the lower sequencing depth, the detected number of mRNA molecules per cell (nCount RNA) was markedly higher in BD Rhapsody data compared to 10X Chromium data (Median nCounts: BD Rhapsody ∼15,500, 10X Chromium: ∼7,350), indicating a subtantially better mRNA capture performance of the microwell-based platform (Figure 2B). We observed this siginficant difference in detected mRNA molecules per cell between the two platforms in individual samples derived from PCa and benign tissues (Figure 2C, Figure S2, and Table S1). Thus, filtering for cells with a defined number of mRNA molecules, markedly reduced the number of high-quality cells predominantly in the 10X Chromium dataset (nCounts > 2000; Figure 2A), indicating superiority of the BD Rhapsody platform in depicting cells with low-mRNA content.

### Prostate cancer tumor microenvironment mapping by microwell- and droplet-based single-cell RNA sequencing

Next, we performed an extensive assessment of the degree of technical biases and the efficacy of the protocols in accurately characterizing distinct cellular populations. Following quality control and filtering procedures, a total of 26,266 PCa and corresponding benign cells (10X Chromium: 16,111 cells; BD Rhapsody: 10,155 cells) were analyzed in terms of their transcriptomes. Using characteristic canonical cell markers, 11 cell clusters were annotated according to the respective cell type (Figure 3A) in the 3 analyzed patients (Figure 3B, individual samples are shown in Figure S4A). We could delineate B cells (*CD79A*), CD4^+^ T cells (*CD4*), CD8^+^ T cells (*CD8*), NK cells (*NKG7*), plasma cells (*JCHAIN*), macrophages & monocytes (*CD68*), mast cells (*KIT*), myofibroblasts (MFB; *ACTA2*), endothelial cells (*CDH5*), basal/intermediate epithelial cells (*KRT5* as well as *KRT19*, *TP63* and low *AR* expression as defined previously in PCa (Chen *et al*, 2021a)), as well as luminal epithelial cells (*KLK3, MSMB*) (Figure 3C and 3D). The cells were derived from both benign prostate and PCa tumor tissues (Figure S4B). Both platforms yielded high-resolution tissue profiles of the analyzed tissues, as illustrated in Figure 3E (see also Figure S4C-F). However, we observed a cell-type-specific bias in cell recovery between the two platforms (Figure 3F), which we investigated in more detail as described further below.

**Figure 3.**
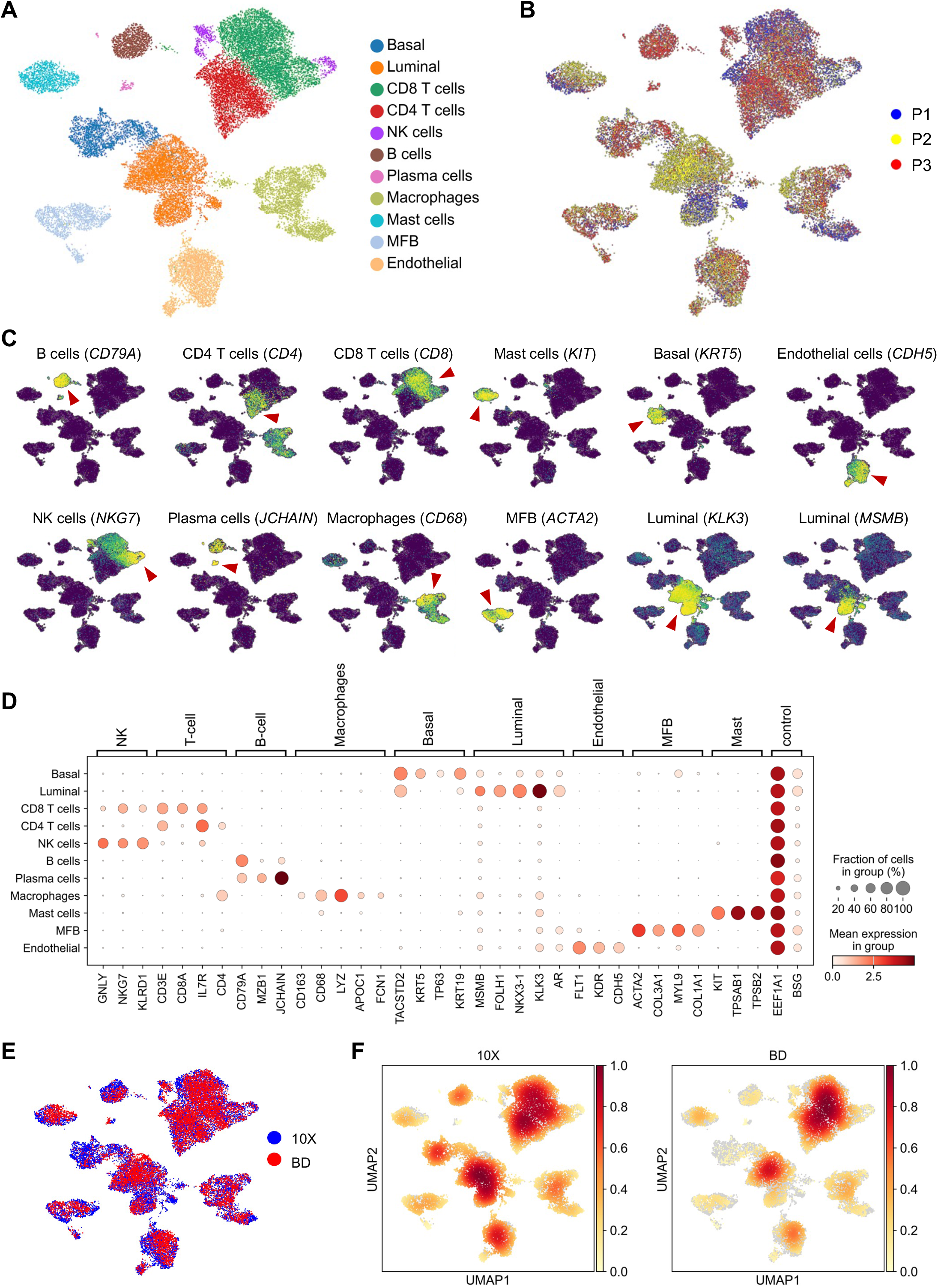
Prostate cancer tumor microenvironment revealed by 10X Chromium and BD Rhapsody. (A) Uniform manifold approximation and projection (UMAP) plot of 26,266 high-quality cells, color-coded by cell type. (B) UMAP plot colored by cells derived from individual patients. (C) UMAP plots colored for the expression of indicated cell-type specific marker genes. (D) Gene expression levels of cell-type specific markers. (E) UMAP plot colored by cells derived from datasets generated with 10X Chromium or BD Rhapsody. (F) UMAP plots showing the cell-density in datasets generated with 10X Chromium or BD Rhapsody. See also **Figure S4**.

### Molecule capture efficiency and sequencing library complexity

The overall higher number of different transcripts detected per cell in the 10X Chromium dataset compared to BD Rhapsody (Figure 2B and 2C) was consistent across individual cell types (Figure 4A). In addition to the previously described differences in sequencing depth, we reasoned that impaired RNA quality/RNA degradation (Figure 2C) might result in less diversity in gene expression due to the degradation of more unstable mRNAs and, subsequently, in lower complexity of transcriptome libraries generated with the BD Rhapsody platform compared to 10X Chromium, respectively. Despite the lower sequencing depth, the median expression level of housekeeping genes (*EEF1A1*, *B2M*, ACTP) was notably higher in the BD Rhapsody dataset both in normalized (Figure 4B) and in raw sequencing data (Figure S5A) in all individual samples (Figure S5B and S5C).

**Figure 4.**
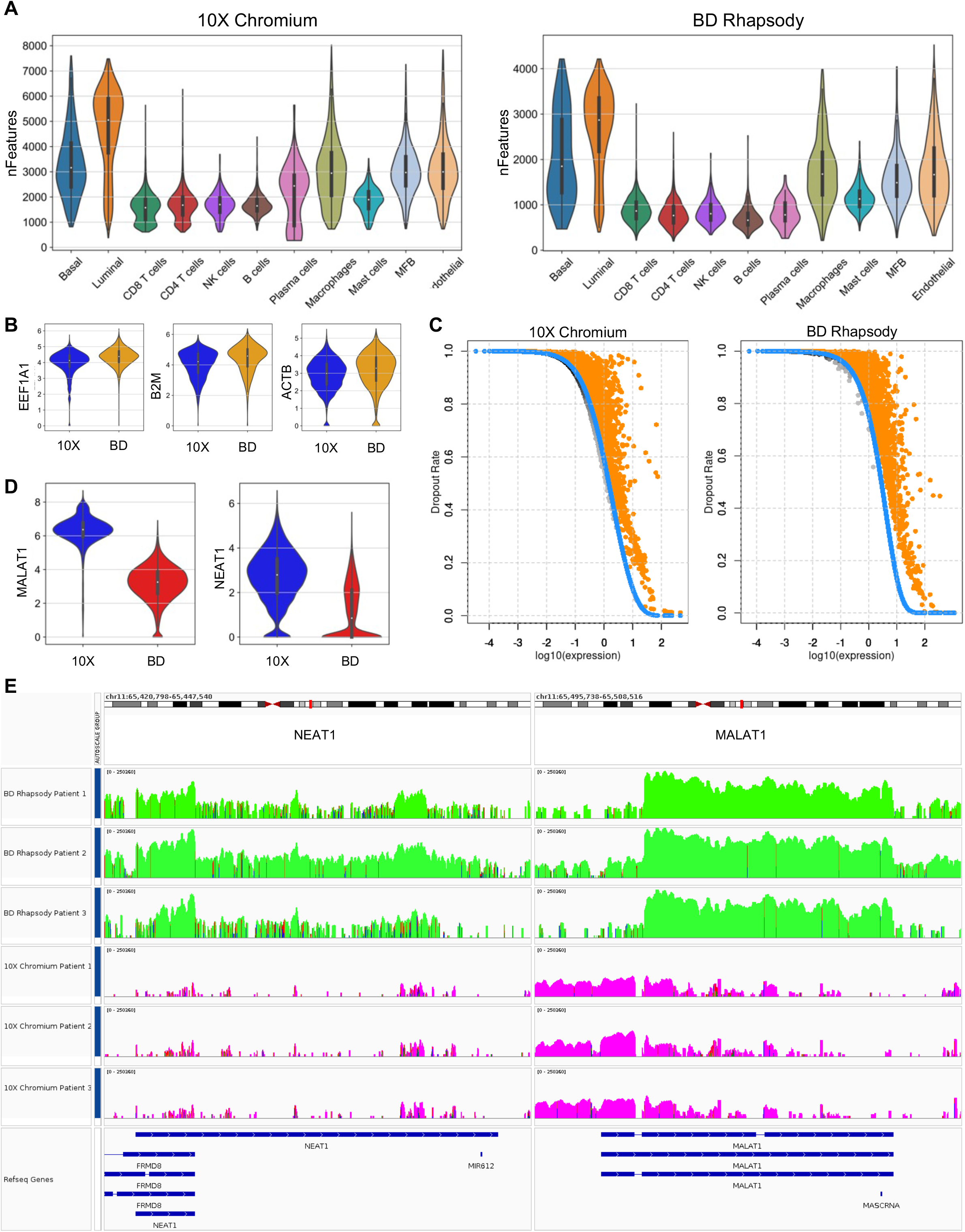
Molecule detection efficiency and sequencing library complexity. (A) Number of genes detected per cell in individual cell types depicted by 10X Chromium and BD Rhapsody. (B) Gene expression levels of indicated house-keeping genes in datasets generated with 10X Chromium and BD Rhapsody. (C) Dropout ratios as a function of log10 expression for 10X Chromium (left) and BD Rhapsody (right). Orange dots represent the significant features under the DANB model (at 1% FDR) while gray dots represent the non-significant features. Blue dots represent the expected dropout probabilities as returned from the DANB model. (D) Gene expression levels of *MALAT1* and *NEAT1* in datasets generated with 10X Chromium and BD Rhapsody. (E) Gene body read coverage of *NEAT1* (left) and *MALAT1* (right). BD Rhapsody samples are represented with green and 10X Chromium samples with purple colours. Data range is set to 0 - 250360 (the min - max read output of both platforms for the specific genomic regions) and a log scale is used to visualize the data and to allow comparisons. The blue and red bands indicate mismatches between the reads and the reference genome’s nucleotide sequences. See also **Figure S5** and **Figure S6**.

Using a modified non-linear Michaelis-Menten equation we were able to correlate the frequency of dropout events (as a measure of dropout ratios) to the gene expression levels. Consistent with prior studies (Mereu *et al*., 2020; Wang *et al*., 2021), genes exhibiting lower expression levels demonstrated higher dropout ratios (Figure 4C). Due to variable library complexity, the dropout probability was found to be higher in the BD Rhapsody dataset (AUC = 5.19) when compared to the 10X Chromium dataset (AUC = 5.05).

Contrary, in the 10X Chromium dataset we found significantly higher expression levels of the Metastasis Associated Lung Adenocarcinoma Transcript 1 (*MALAT1*, also known as *NEAT2*) and the Nuclear-Enriched Abundant Transcript 1 (*NEAT1*; Figure 4D). This marked difference in *MALAT1* and *NEAT1* expression between the two platforms was visible in all individual cell types of the prostate TME (Figure S6A). Concordantly, our recently published Non-Small Cell Lung Cancer (NSCLC) scRNA-seq atlas (Salcher *et al*., 2022) revealed that *MALAT1* and *NEAT1* are highly abundant in various cells of the lung cancer TME (Figure S6B) and noticeable higher expressed in datasets generated with 10X Chromium compared to BD Rhapsody (Figure S6C), respectively. *MALAT* and *NEAT1* are both nuclear retained long non-coding RNAs (lncRNAs). *MALAT* lacks a poly(A) tail (Wilusz *et al*, 2012), whereas NEAT1 does undergo some degree of polyadenylation, which primarily applies to the NEAT1_1 isoform (Naveed *et al*, 2021).

To further investigate the difference in the expression of *MALAT1* and *NEAT1* between the two platforms, we determined the gene body coverage using the IGV viewer (version 2.16.1) (Figure 4E). BD Rhapsody exhibited an increased number of reads mapping to the gene body of both genes, which was exponentially higher than 10X Chromium. In the BD Rhapsody data, the reads mapped unevenly on the gene body of *MALAT1*, more specifically there appears to be a bias towards the genomic regions 65,498,957 - 65,506,516 with very few reads mapping to exon 1 of *MALAT1* (RefSeq gene: NR_144568.1). In contrast, in the 10X Chromium data, the bias appears to be towards exon 1 with the majority of reads mapping on the genomic region 65,497,738 - 65,502,629 of Chromosome 11. These marked differences were visible in all individual patients. The distribution of reads towards exon 1 seems to capture the expression of *MALAT1* with higher sensitivity, which may explain the observed difference in *MALAT1* expression between the two platforms. In the case of *NEAT1*, the gene coverage followed a similar pattern in both platforms. More specifically most of the reads mapped on the genomic regions 65,422,798 - 65,426,543 and 65438091 - 65,445,540 of Chromosome 11. It is important to note that the genomic region 65,422,798 - 65,426,543 of Chromosome 11 overlaps with exon 13 of the FRMD8 gene (RefSeq gene: XR_007062512.1), which may lead to the reads appearing as “multi-mappers” in the downstream analysis. Of note, in the case of BD Rhapsody, there appears to be an increased percentage of mismatches in the nucleotide sequences of the reads mapping to the reference genome of *NEAT1*, which may possibly affect the gene counts in the downstream analysis.

We next conducted a comparative analysis of the top 200 differentially expressed genes (DEGs) exemplarily in luminal epithelial cells and in endothelial cells, as identified in PCa and benign prostate tissues by 10X Chromium and BD Rhapsody (Figure 5A). We observed a partially overlapping expression pattern of DEGs in both, luminal epithelial (139/200 overlapping DEGs) and endothelial cells (145/200 overlapping DEGs), between both platforms (Figure 5B). To further assess the suitability of the protocols for characterizing cell types, we evaluated their sensitivity in detecting population-specific expression markers and uncovered a significant disparity in their ability to discern marker genes. The prostate-specific-antigen (PSA, *KLK3*) as well as other prostate epithelial markers such as *KLK2*, *KLK4*, or *ACPP* were highly expressed in luminal epithelial cells in data generated with 10X Chromium as well as BD Rhapsody (Figure 5C). However, particularly endothelial cell population markers were detected with varying sensitivity (Figure 5C, Table S2). Both, the interferon alpha-inducible protein 27 (*IFI27*), previously described as marker for capillary endothelial cells of the human lung (Schupp *et al*, 2021), as well as the key endothelial cell marker von-Willebrand factor (*VWF*), exhibited markedly higher expression in the 10X Chromium dataset compared to the BD Rhapsody dataset, respectively (Figure 5C). One explanation for this discrepancy is that specific RNAs that are more sensitive to RNA degradation are lost during the BD Rhapsody protocol. However, stability of mRNAs is influenced by various factors, such as the presence of stabilizing or destabilizing elements within the mRNA sequence, interactions with RNA-binding proteins, and the cellular context. The current literature does not provide clear evidence to support the notion that these particular mRNAs are inherently unstable or rapidly degraded. Conversely, we observed a trend towards elevated expression levels of *CD34* (a well-known marker of progenitor cells of blood vessels in endothelial cells depicted with the BD Rhapsody platform (Figure 5C). The discrepancies in depicting endothelial cell population markers were visible in the raw sequencing data (Figure S7A) as well as in the filtered and normalized data (Figure 5D) in all individual samples (Figure 5E). In concordance with the overall elevated expression levels of lncRNAs (Figure 4D), the expression of *MALAT1* and *NEAT1* was also significantly higher in endothelial cells depicted with the10X Chromium protocol (Figure S7B, Table S2).

**Figure 5.**
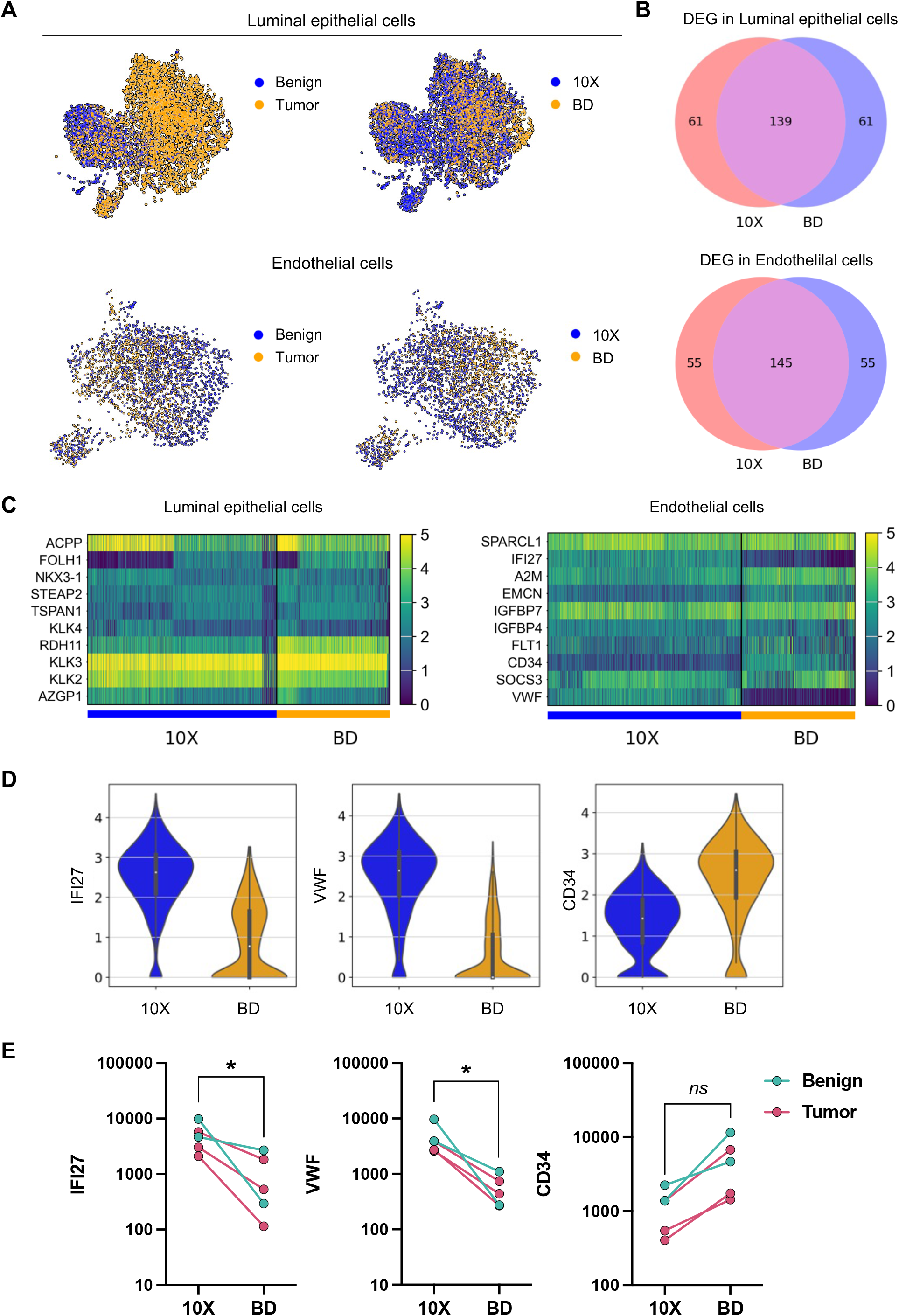
Platform-specific gene quantification and cell-type marker identification. (A) UMAP blots of luminal epithelial cells and endothelial cells colored by cells derived from benign and PCa tumor tissues as well as by cells derived from datasets generated with 10X Chromium and BD Rhapsody. (B) VENN diagram of top 200 DEG detected by 10X Chromium or BD Rhapsody in luminal epithelial cells (upper panel) and endothelial cells (lower panel). (C) Top expressed genes in luminal epithelial cells and endothelial cells in datasets generated with 10X Chromium and BD Rhapsody. (D) Gene expression levels of *IFI27*, *VWF*, and *CD34* detected in endothelial cells by 10X Chromium and BD Rhapsody. (E) Gene expression levels of *IFI27*, *VWF*, and *CD34* in individual samples. Each dot refers to a sample (benign or tumor tissue) with at least 40 endothelial cells in both 10X Chromium and BD Rhapsody groups. Paired t-test, *p ≤ 0.05. See also **Figure S7**.

In summary, the comparison of the 10X Chromium and BD Rhapsody platforms revealed similarities in detecting differentially expressed genes (DEGs), while also exhibiting platform-dependent variability in detecting specific RNAs, including lncRNAs and cell-type specific population markers.

### Platform-dependent cellular composition in single-cell RNA sequencing data

The overall lower mRNA count in the 10X Chromium dataset (Figure 2B and 2C) may particularly affect detection and characterization of cells with low-mRNA content, such as various immune cells. Concordantly, we observed that CD4^+^ T cells, CD8^+^ T cells, and to a lesser extend NK cells were primarily affected by filtering for cells above a minimum mRNA amount in the 10X Chromium dataset (Figure 6A).

**Figure 6.**
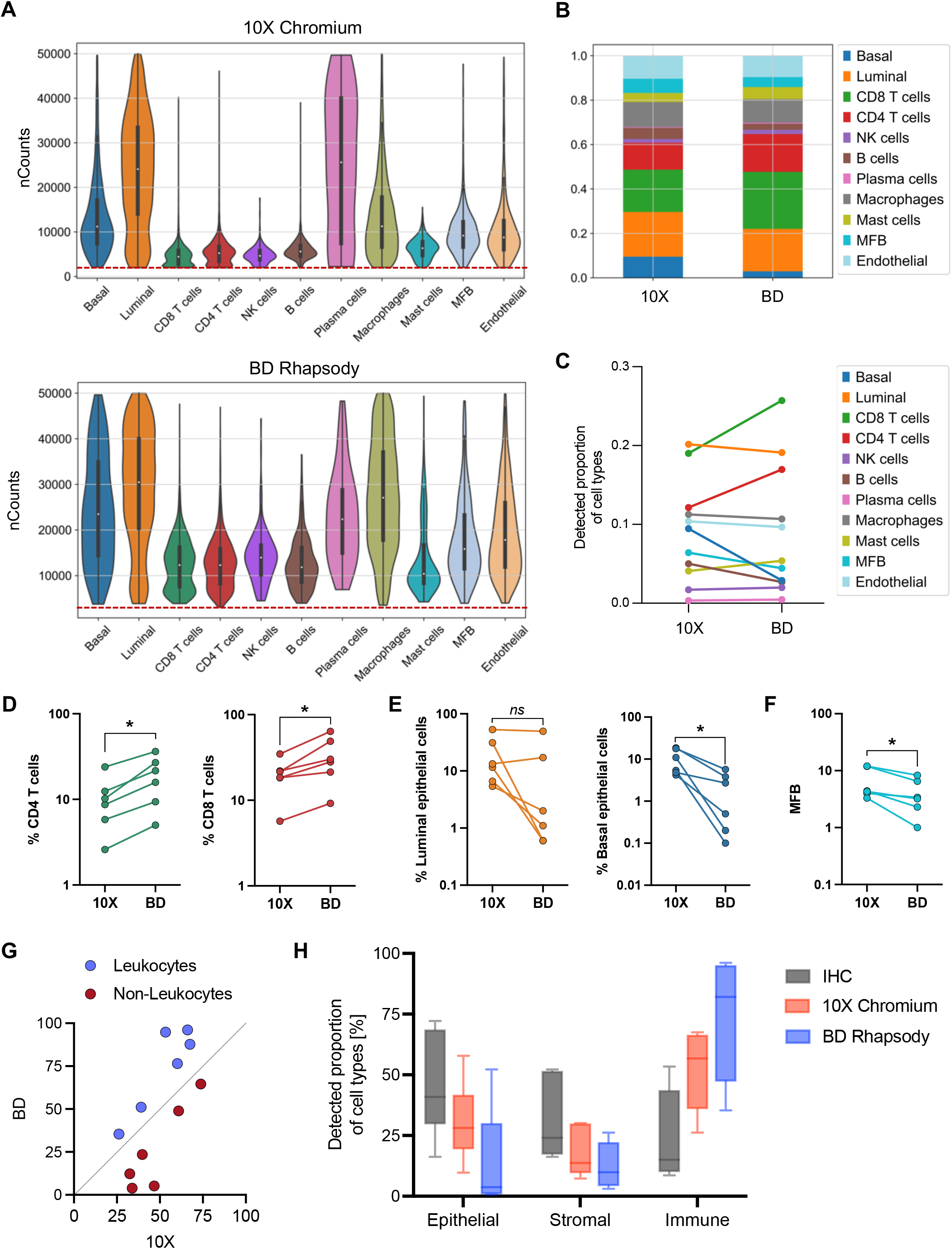
Platform-dependent cellular composition in single-cell RNA sequencing data. (A) Number of captured transcripts (total counts) in individual cell types recovered with 10X Chromium (upper panel) and BD Rhapsody (lower panel). (B) Relative cell-type composition in benign prostate and PCa tissues from three individual patients in data generated with 10X Chromium and BD Rhapsody. (C) Detected proportion of cell-types in benign prostate and PCa tissues from three individual patients in data generated with 10X Chromium vs. BD Rhapsody. (D-F) The proportion of CD4 T cells and CD8 T cells (D), luminal and basal epithelial cells (E), and myofibroblasts (MFB) (F), depicted by 10X Chromium vs. BD Rhapsody in individual samples. Paired t-test, *p ≤ 0.05. (G) The proportion of leukocytes and non-leukocytes depicted by 10X Chromium vs BD Rhapsody. (H) Proportions of epithelial, stromal, and CD45^+^ immune cells in benign prostate and PCa tissues from three individual patients determined by IHC and by scRNA-seq using 10X Chromium or BD Rhapsody. See also **Figure S8**.

Consequently, CD4^+^ T cells and CD8^+^ T cells were detected at significantly smaller proportions by 10X Chromium compared to BD Rhapsody (Figure 6B and 6C), in all individual samples (Figure 6D). Additionally, we noted a trend towards higher NK cell numbers in the BD Rhapsody dataset (Figure S8A). Conversely, epithelial cells (Figure 6E) and myofibroblasts (Figure 6F), were substantially better represented in the 10X Chromium dataset. Particularly basal epithelial cells were significantly better depicted by the 10X Chromium platform in all six analyzed samples (Figure 6E). Primary prostate epithelial cells are highly sensitive to anoikis and cell stress, which may lead to their loss. Consequently, we observe a substantial increase in %MT expression in basal epithelial cells compared to other cell types in the BD Rhapsody dataset, which might at least in part reduce the cell number after filtering (Figure S8B). Overall, the leukocyte fraction was particularly represented in BD Rhapsody data, whereas non-leukocytes, including epithelial cells and myofibroblasts were depicted at higher proportions by 10X Chromium (Figure 6G).

PCa is generally deemed as immunologically ‘cold’ tumor with relatively low immune cell infiltration, as also demonstrated by IHC analysis showing a higher abundance of epithelial and stromal cells in all analyzed benign prostate and PCa tumor tissues compared to CD45+ leukocytes (Figure 6H). Contrary, both scRNA-seq protocols resulted in an overrepresentation of immune cells and an underrepresentation of the epithelial/stromal compartment. This phenomenon may be mostly attributable to the process of tissue dissociation, nonetheless, the composite data propose that the microwell-based scRNA-seq technique preferentially enriches for immune cells, while the droplet-based protocol recovers epithelial/stromal cells at comparatively elevated proportions (Figure 6H).

## DISCUSSION

We systematically compared two of the most broadly used high-throughput scRNA-seq technologies (i.e. 10X Chromium and BD Rhapsody) using paired samples from surgically resected PCa and the respective healthy prostate tissues. Contrary to previous comparative analysis (Chen *et al*., 2021b; Colino-Sanguino *et al*., 2023; Gao *et al*., 2020; Mereu *et al*., 2020; Natarajan *et al*., 2019; Wang *et al*., 2021; Yamawaki *et al*., 2021; Zhang *et al*., 2019b; Ziegenhain *et al*., 2017), we simultaneously processed a sufficient number of samples derived from complex tissues for a statistically sound comparison. Our analysis revealed differences between the two platforms in converting RNA molecules into sequencing libraries, impacting the reliability and accuracy of the resulting scRNA-seq data. This information is essential for scientists asking for the right technology platform for their experimental research question: which platform is indeed appropriate to reliably detect expression profiles of single cells for the posed question.

First, our comparative analysis of QC metrics revealed that the droplet-based 10X Chromium platform yields a higher effective cell capture rate and allows a more sensitive detection of genes per cell, while the microwell-based BD Rhapsody protocol exhibits a substantially higher number of mRNA molecules detected per cell. We demonstrate that the BD Rhapsody platform captures a higher proportion of mitochondrial transcripts, typically associated with increased cell stress. However, we found no evidence of procedural stress as a contributing factor. One possible explanation for the elevated proportion of mitochondrial transcripts in the BD Rhapsody dataset could be a more efficient disruption of organelle membranes during the cell lysis step compared to the 10X Chromium protocol, which employs a markedly lengthier yet potentially weaker cell lysis procedure.

Second, we demonstrate that the sample-multiplexing capability in the BD Rhapsody workflow using sample-tags, markedly impairs RNA quality in freshly isolated single-cell suspensions derived from complex tissues. On the plus side, sample-multiplexing reduces technical bias caused by batch effects and substantially lowers sequencing library preparation costs. However, as RNA quality is a critical determinant of scRNA-seq data reliability and accuracy, the BD Rhapsody sample-tag staining procedure might compromise specific mRNAs, impair sequencing library complexity, and thus potentially affect the outcome of the scRNA-seq experiment.

Third, we report that varying library complexities and the observed transcript-specific mRNA capture efficacy of both protocols affect their ability to quantify gene expression levels and identify cell-type markers. Overall, the BD Rhapsody protocol demonstrated a higher gene dropout probability. The relative high abundance of mitochondrial and house-keeping transcripts in the BD Rhapsody data may indirectly contribute to elevated dropouts of genes exhibiting lower expression levels.

Although the vast majority of transcripts shows congruent expression pattern in different cell types with both platforms, our analysis also revealed substantial differences for certain genes, exemplarily shown for *IFI27*, *VWF* or *CD34*, putative marker genes of prostate endothelial cells. In turn, this discrepancy in amplifying certain types of RNA molecules, might significantly impact the underlying composition of depicted tissue profiles and affect the exploratory value of generated scRNA-seq datasets. Of note, we observed remarkably high expression levels of the lncRNAs *MALAT1* and *NEAT1* exclusively in 10X Chromium data. We could confirm this platform-specific differences in depicting lncRNAs in our recently published NSCLC scRNA-seq atlas. Gene coverage analysis revealed a pronounced 3’ end bias in BD Rhapsody read distribution for *MALAT1*, whereas the 10X Chromium platform exhibited a distinct bias towards exon 1 at the 5’ end of the gene. Thus, we speculate that a higher sensitivity of exon 1 read capture may account for the observed variation in *MALAT1* expression between the two platforms. Overall, we advocate to consider potential platform-mediated gene expression bias when interpreting data or comparing results derived from different scRNA-seq protocols.

Finally, our direct comparative analysis corroborates that the BD Rhapsody protocol excels in characterizing cells with low-mRNA content, capturing leukocytes more effectively. This important benefit of the microwell-based platform is based on a significantly better mRNA capture efficiency. In line with our recent observation in lung cancer (Salcher *et al.,* 2022), the detected number of mRNA molecules per cell was markedly higher in BD Rhapsody data compared to 10X Chromium, respectively. Conversely, the droplet-based protocol performed noticeably better in depicting epithelial cells, particularly prostate basal epithelial cells, as well as myofibroblasts. Considering that the microwell-based BD Rhapsody platform is validated for cells smaller than 20 µm, we hypothesize that the loss of epithelial cells or myofibroblasts may be attributable to their larger size relative to leukocytes. Hence, processing single-cell suspensions derived from complex tissues using microwell-based platforms might result in a bead-exclusion phenomenon that results in loss of larger cell types. In accordance, our recent scRNA-seq studies on liver tissues processed with the BD Rhapsody platform, resulted in an underrepresentation of hepatocytes (typical size > 20 µM), the primary liver cells (Hautz *et al*., 2023). Overall, our findings propose that the choice of the scRNA-seq protocol might substantially influence the composition of captured cell types, and thus researchers must carefully evaluate their research questions before selecting the most appropriate platform.

While our results provide valuable insights into the performance of these scRNA-seq protocols, it is important to acknowledge several potential limitations of our study including platform-specific variables in handling, processing, and library preparation. Future studies with more diverse samples could further enhance our understanding of scRNA-seq platform performance. It is also important to note, that optimizing the workflow might improve the performance of both platforms. Lastly, researchers should account for potential biases during data analysis and ensure their methods suit the specific platform and research question. Despite these potential limitations, our findings offer valuable insights into the performance characteristics of the 10X Chromium and BD Rhapsody platforms in the context of scRNA-seq analysis of prostate cancer tissues.

In conclusion, our study emphasizes the importance of considering the distinct characteristics of scRNA-seq technologies. Researchers must carefully evaluate their research questions and select the most appropriate platform accordingly. Furthermore, we advocate to be cautious when comparing data generated by different platforms, as platform-dependent variability in detecting population-specific markers and distinctive cell subpopulations may lead to discrepancies in the interpretation of the results. In that direction, our results might lead to informed algorithms for better data integration of different platforms. Future research is necessary to elucidate the underlying reasons for these differences and to optimize scRNA-seq protocols for obtaining reliable and accurate data in various biological contexts.

## MATERIALS AND METHODS

### Experimental model and subject details

#### Human subjects

The local ethics committee (EK no. 1017/2018; 1072/2018) approved the use of tissue samples obtained from fresh radical prostatectomy (RP) specimens. Written informed consent is available from all patients. As described by our group, the malignancy or benignity of the tissue was confirmed within 1 h after surgery. Subsequently, tissue dissociation and FACS sorting of freshly isolated cells was performed to obtain a FACS-sorted PCa and corresponding benign prostate single-cell suspension (Heidegger *et al*., 2022).

Samples of NSCLC tumor tissues and matched benign lung tissues were obtained from surgical specimens of patients undergoing resection at the Department of Visceral, Transplant and Thoracic Surgery (VTT), Medical University Innsbruck, Austria, and in collaboration with the INNPATH GmbH, Innsbruck, Austria, after obtaining informed consent in accordance with a protocol reviewed and approved by the Institutional Review Board at the Medical University Innsbruck, Austria (study code: AN214-0293 342/4.5).

### Reagents and Tools Table

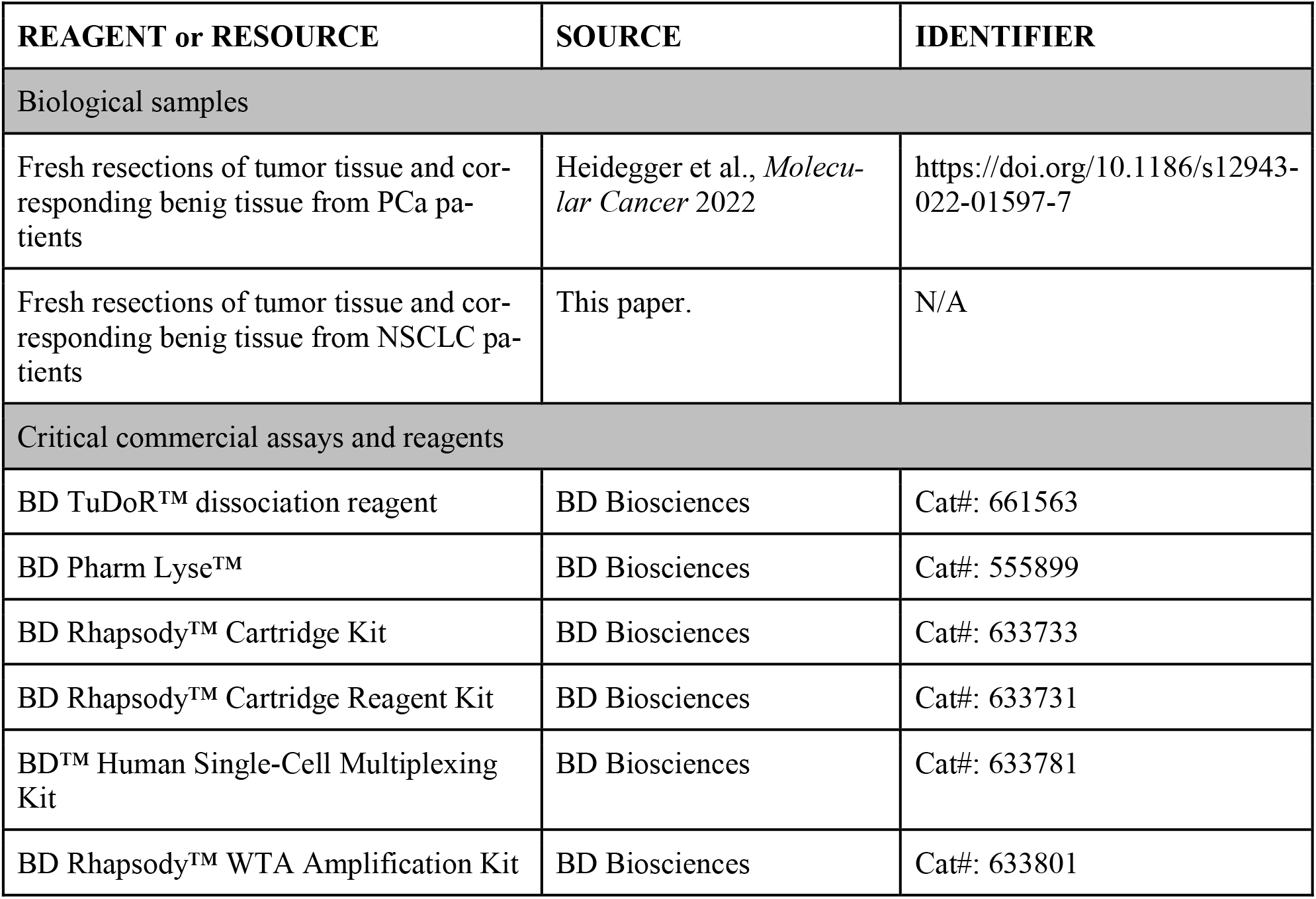
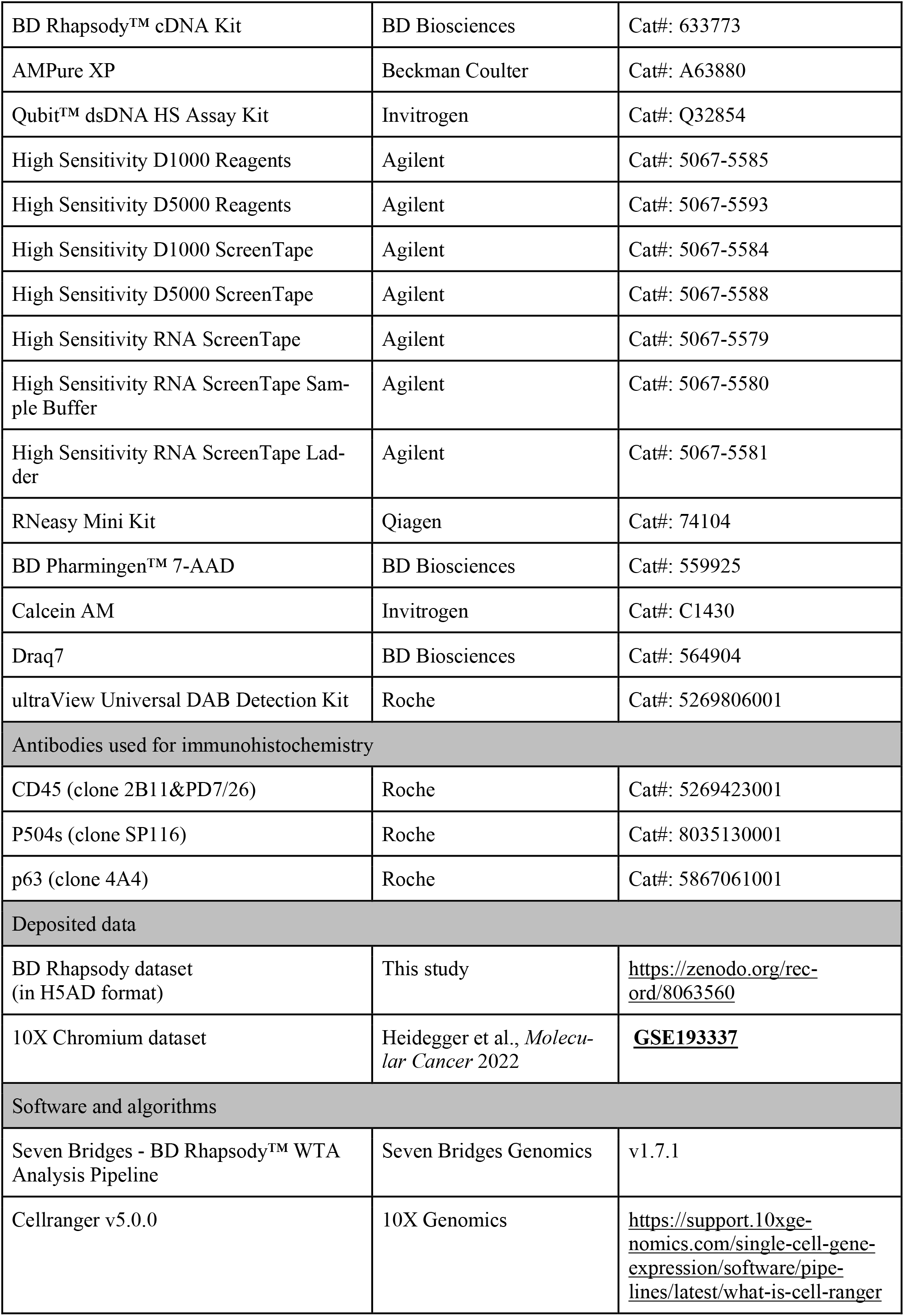
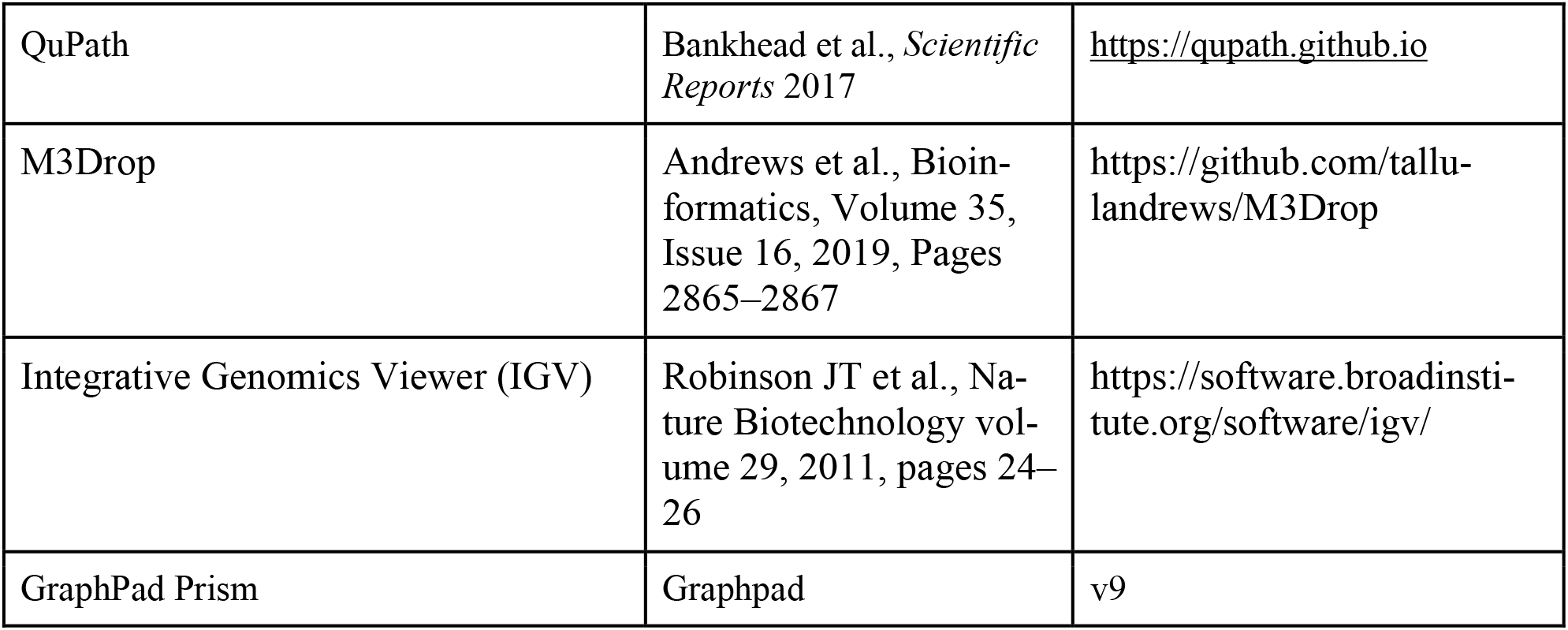

### Methods and Protocols

#### 10X Chromium library preparation and sequencing

The droplet-based 10X Chromium system employs the GemCode technology, which utilizes gel beads in emulsion to achieve barcoding *(Eisenstein, 2015)*. This involves combining a suspension of single cells with gel beads containing barcoded oligonucleotides and reagents for reverse transcriptase (RT) in an oil environment, resulting in the formation of nanoliter-scale droplets that facilitate cDNA synthesis. Subsequently, the droplets are pooled, dissolved, and used to create a sequencing library containing unique molecular identifiers (UMIs). The microfluidic chips used in this system can accommodate up to eight samples simultaneously, each containing up to 10,000 cells. Samples obtained from four PCa patients were converted to scRNA-seq libraries using the Chromium Next GEM Single-Cell 3’Kit v3.1 (10X Genomics) as described previously (Heidegger *et al*., 2022). Three out of these four patients (∼48,000 cells) were processed in parallel with the BD Rhapsody platform.

#### BD Rhapsody library preparation and sequencing

The BD Rhapsody platform uses a single cartridge with more than 200,000 microwells, wherein up to 30,000 individual cells are isolated together with UMI-barcoded magnetic mRNA capture-beads. Upon cell lysis the mRNAs are captured and retrieved together with the beads out of the microwells. All beads are pooled, and subsequently the RT step is performed within a single tube. Freshly isolated, FACS-sorted single-cells were immediately processed with the BD Rhapsody scRNA-seq platform (BD Biosciences). The BD Single-Cell Multiplexing Kit (BD Biosciences) was used to combine and load two samples (tumor tissue and corresponding benign tissue) onto a single BD Rhapsody cartridge (BD Biosciences). Sample-tag staining was performed according to the manufacturer’s protocol (sample-tag staining at room temperature for 20 minutes and washing by centrifugation at 400 g for 5 minutes). Single-cell isolation in microwells (cell load: 20 minutes incubation at room temperature) with subsequent cell-lysis and capturing of poly-adenylated mRNA molecules with barcoded, magnetic captured-beads was performed according to the manufacturer’s instructions. Beads were magnetically retrieved from the microwells, pooled into a single tube before reverse transcription. Unique molecular identifiers (UMIs) were added to the cDNA molecules during cDNA synthesis. Whole transcriptome amplification (WTA) and sample-tag sequencing libraries were generated according to the BD Rhapsody single-cell whole-transcriptome amplification workflow. The quantity and quality of the sequencing libraries was analyzed with the Qubit dsDNA HS (High Sensitivity) assay kit (Invitrogen) and the 4200 TapeStation (Agilent) system. Libraries were sequenced on the Novaseq 6000 system (Illumina) targeting a sequencing depth of 45.000 reads/cell.

#### Preprocessing and quality control of scRNA-seq data

Bioinformatic pre-processing of the obtained FastQ sequencing files was performed via the cloud-based Seven Bridges Platform environment (Seven Bridges Genomics) using the BD Rhapsody WTA Analysis Pipeline app making use of the sample-tag workflow to achieve csv-files with cell codes and gene list for each individual sample. Fastq sequencing data derived from 10X Chromium were mapped to the human genome (build GRCh38) using the CellRanger software (10X Genomics, v3.1) to achieve mtx, barcodes and genes files for each sample as described previously by our group (Heidegger *et al*., 2022). Sample files were imported in Scanpy version 1.9.1 running with Python version 3.8 (Wolf *et al*, 2018) and loaded into AnnData (Virshup *et al*, 2021) for further processing with scverse tools to perform removal of empty cell beats and cell barcodes without sufficient captured transcripts (>100 genes, gene occurance in >3 cells). Thereafter, all samples were imported into a single anndata object (h5ad) by the concatenation “outer” join function to maintain all platform/sample specific genes since objects had differing variables. Empty genes (variables) for each sample were filled up with 0 values.

#### Integration of scRNA-seq datasets

Quality control was performed using scanpy, only retaining cells with (I) between 200 and 8000 detected genes, (II) between 1000 and 50,000 transcripts, and (III) <30% mitochondrial transcripts. The 2000 most highly variable genes (HVGs) were selected using scanpy’s highly_variable_genes function with the options flavor = ”seurat_v3” and batch_key = ”sample”. Cell transcriptomes were embedded into a batch-corrected low-dimensional latent space using scVI (Gayoso *et al*, 2022; Xu *et al*, 2021) treating each sample as a separate batch. A neighborhood graph and uniform manifold approximation and projection (UMAP) embedding (Becht *et al*, 2018) were computed based on the scVI latent space. Cell types were annotated based on unsupervised clustering with the Leiden algorithm (Traag *et al*, 2019) and known marker genes specific for epithelial cells, endothelial cells, myofibroblasts and immune cell types.

#### Differential gene expression testing

For each cell type and patient, we summed up transcript counts for each gene that is expressed in at least 25% of cells and at least in 3 samples using decoupler-py (Badia *et al*, 2022). Pseudo-bulk samples consisting of fewer than 1000 counts or 40 cells were discarded. Pseudo-bulked data for endothelial and luminal epithelial cells were used for further differential gene expression testing between the two different platforms using DESeq2 (Love *et al*, 2014) which has been demonstrated to perform well (Squair *et al*, 2021).

#### Dropout ratio analysis

The dropout ratios were correlated to gene expression levels by fitting a modified non-linear Michaelis-Menten equation introduced in the M3Drop R package (Andrews & Hemberg, 2019). To account for zeros resulting from insufficient sequencing depth, we used the depth-adjusted negative binomial model (DANM). DANB assumes that each observation follows a negative binomial distribution, where the mean is proportional to both the mean expression of the specific gene and the relative sequencing depth of the corresponding cell.

#### Gene body coverage analysis

In order to visualize the gene body coverage, the BAM files produced in the pre-processing step were directly loaded in the IGV viewer (Robinson *et al*, 2011) (version 2.16.1) as different tracks. The viewing panel was centered around the *NEAT1* and *MALAT1* genomic regions. To account for the difference in sequencing depth the maximum data threshold was set to 250,360 (which is the maximum number of reads found to map on the genomic regions of *NEAT1* and *MALAT1*) and the data were log transformed for comparison reasons.

#### RNA quality of single-cell suspensions derived from NSCLC and normal lung tissue

Surgically resected NSCLC tumor tissues and corresponding benign lung tissues were minced into small pieces (< 1 mm) on ice and enzymatically digested with agitation for 30 minutes at 37°C using the BD TuDoR™ dissociation reagent (BD Biosciences). The obtained single-cell solution was sieved through a 70 µM cell strainer (Corning) and red blood cells were removed using the BD Pharm Lyse™ lysing solution (BD Biosciences). Cells were counted and viability assessed with the BD Rhapsody scRNA-seq platform (BD Biosciences) using Calcein-AM (Invitrogen) and Draq7 (BD Biosciences). Immediately, >1×10^6^ cells of the obtained single-cell suspensions were subjected to the sample-tag staining procedure (20 min RT, 3x washing by 5 min centrifugation at 400 rpm). Total RNA was isolated before (T1) and after (T2) the sample-tag staining procedure using the RNeasy Mini kit (Qiagen) and RNA quality (RNA integrity number, RIN) was assessed with the High Sensitivity RNA ScreenTape assay (Agilent) and the 4200 TapeStation (Agilent) system according to the manufacturer’s instructions.

#### Immunohistochemistry

Immunohistochemical analysis with slides from the investigated 3 PCa patients was executed. Locations of interest on the slides were chosen based on slides from the same blocks stained with the markers P504s (clone SP116, Roche, #8035130001) and p63 (clone 4A4, Roche, #5867061001) to identify tumor regions or benign prostate regions. CD45 was stained (clone 2B11&PD7/26, Roche, #5269423001) to identify immune cells. Antibody staining was detected with the ultraView Universal DAB Detection Kit (Roche, #5269806001). The slides were scanned with Hamamatsu NanoZoomer S210 (Hamamatsu, Shizuoka, Japan) with a 40x magnification. Representative regions were selected by an expert prostate pathologist (G.S.) and the proportions of stromal, epithelial, and CD45^+^ cells were calculated using the QuPath software for digital pathology image analysis (Bankhead *et al*, 2017).

#### Quantification and statistical analysis

Statistical analysis was performed GraphPad Prism. Single cell-data were aggregated into pseudo-bulk samples by biological replicates. Significance levels on the statistical tests are indicated in the figure captions.

## ACKNOWLEDGMENTS

This work was supported by the Austrian Science Fund FWF (Grant-No. TAI-697) (DW), the “In Memoriam Gabriel Salzner Stiftung” (DW), FFG grant Austrian Research Promotion Agency, 858057 (HD FACS, SSo)., TWF (Grant-No F.16733/5-2019) (IH).

## AUTHOR CONTRIBUTIONS

Conceptualization, StS, GU, DW, SiS, and AP; data analysis, GU, StS, GF, SiS; human samples, IH, AP, DW; single-cell sequencing, StS, AN, AK; sample analysis, StS; orthogonal validation, GS, CS, IH; visualization, StS, AS; writing-original draft, StS; writing - review and editing, all authors; funding acquisition: DW, SiS, IH.

## DATA AVAILABILITY

Processed scRNA-seq data from this study has been deposited on Zenodo (https://zenodo.org/record/8063560) as listed in the key resources table. Raw data is not made available due to privacy concerns.

## SUPPLEMENTAL INFORMATION

**Supplemental Table 1: Quality metrics of individual samples, related to Figure 1**.

**Supplemental Table 2: Gene expression in endothelial cells in individual samples, related to Figure 5**.

## REFERENCES

Andrews TS, Hemberg M (2019) M3Drop: dropout-based feature selection for scRNASeq. Bioinformatics (Oxford, England) 35: 2865–2867

Azizi E, Carr AJ, Plitas G, Cornish AE, Konopacki C, Prabhakaran S, Nainys J, Wu K, Kiseliovas V, Setty M et al (2018) Single-Cell Map of Diverse Immune Phenotypes in the Breast Tumor Microenvironment. Cell 174: 1293–1308 e1236

Badia IMP, Velez Santiago J, Braunger J, Geiss C, Dimitrov D, Muller-Dott S, Taus P, Dugourd A, Holland CH, Ramirez Flores RO et al (2022) decoupleR: ensemble of computational methods to infer biological activities from omics data. Bioinform Adv 2: vbac016

Bankhead P, Loughrey MB, Fernandez JA, Dombrowski Y, McArt DG, Dunne PD, McQuaid S, Gray RT, Murray LJ, Coleman HG et al (2017) QuPath: Open source software for digital pathology image analysis. Sci Rep 7: 16878

Becht E, McInnes L, Healy J, Dutertre CA, Kwok IWH, Ng LG, Ginhoux F, Newell EW (2018) Dimensionality reduction for visualizing single-cell data using UMAP. Nat Biotechnol

Chen S, Zhu G, Yang Y, Wang F, Xiao YT, Zhang N, Bian X, Zhu Y, Yu Y, Liu F et al (2021a) Single-cell analysis reveals transcriptomic remodellings in distinct cell types that contribute to human prostate cancer progression. Nature cell biology 23: 87–98

Chen W, Zhao Y, Chen X, Yang Z, Xu X, Bi Y, Chen V, Li J, Choi H, Ernest B et al (2021b) A multicenter study benchmarking single-cell RNA sequencing technologies using reference samples. Nat Biotechnol 39: 1103–1114

Colino-Sanguino Y, Fuente LRdl, Gloss B, Law AMK, Handler K, Pajic M, Salomon R, Gallego-Ortega D, Valdes-Mora F (2023) Systematic comparison of high throughput Single-Cell RNA-Seq platforms in complex tissues. bioRxiv: 2023.2004.2004.535585

Eisenstein M (2015) Startups use short-read data to expand long-read sequencing market. Nat Biotechnol 33: 433–435

Fan HC, Fu GK, Fodor SP (2015) Expression profiling. Combinatorial labeling of single cells for gene expression cytometry. *Science (New York*, NY) 347: 1258367

Gao C, Zhang M, Chen L (2020) The Comparison of Two Single-cell Sequencing Platforms: BD Rhapsody and 10x Genomics Chromium. Curr Genomics 21: 602–609

Gayoso A, Lopez R, Xing G, Boyeau P, Valiollah Pour Amiri V, Hong J, Wu K, Jayasuriya M, Mehlman E, Langevin M et al (2022) A Python library for probabilistic analysis of single-cell omics data. Nat Biotechnol 40: 163–166

Gierahn TM, Wadsworth MH, 2nd, Hughes TK, Bryson BD, Butler A, Satija R, Fortune S, Love JC, Shalek AK (2017) Seq-Well: portable, low-cost RNA sequencing of single cells at high throughput. Nat Methods 14: 395–398

Goveia J, Rohlenova K, Taverna F, Treps L, Conradi LC, Pircher A, Geldhof V, de Rooij L, Kalucka J, Sokol L et al (2020a) An Integrated Gene Expression Landscape Profiling Approach to Identify Lung Tumor Endothelial Cell Heterogeneity and Angiogenic Candidates. Cancer cell 37: 421

Goveia J, Rohlenova K, Taverna F, Treps L, Conradi LC, Pircher A, Geldhof V, de Rooij L, Kalucka J, Sokol L et al (2020b) An Integrated Gene Expression Landscape Profiling Approach to Identify Lung Tumor Endothelial Cell Heterogeneity and Angiogenic Candidates. Cancer cell 37: 21–36 e13

Guo X, Zhang Y, Zheng L, Zheng C, Song J, Zhang Q, Kang B, Liu Z, Jin L, Xing R et al (2018) Global characterization of T cells in non-small-cell lung cancer by single-cell sequencing. Nature medicine 24: 978–985

Han X, Wang R, Zhou Y, Fei L, Sun H, Lai S, Saadatpour A, Zhou Z, Chen H, Ye F et al (2018) Mapping the Mouse Cell Atlas by Microwell-Seq. Cell 173: 1307

Hautz T, Salcher S, Fodor M, Sturm G, Ebner S, Mair A, Trebo M, Untergasser G, Sopper S, Cardini B et al (2023) Immune cell dynamics deconvoluted by single-cell RNA sequencing in normothermic machine perfusion of the liver. Nature communications 14: 2285

Hedlund E, Deng Q (2018) Single-cell RNA sequencing: Technical advancements and biological applications. Molecular aspects of medicine 59: 36–46

Heidegger I, Fotakis G, Offermann A, Goveia J, Daum S, Salcher S, Noureen A, Timmer-Bosscha H, Schafer G, Walenkamp A et al (2022) Comprehensive characterization of the prostate tumor microenvironment identifies CXCR4/CXCL12 crosstalk as a novel antiangiogenic therapeutic target in prostate cancer. Molecular cancer 21: 132

Hwang B, Lee JH, Bang D (2018) Single-cell RNA sequencing technologies and bioinformatics pipelines. Exp Mol Med 50: 1–14

Ilicic T, Kim JK, Kolodziejczyk AA, Bagger FO, McCarthy DJ, Marioni JC, Teichmann SA (2016) Classification of low quality cells from single-cell RNA-seq data. Genome Biol 17: 29

Ji F, Sadreyev RI (2019) Single-Cell RNA-seq: Introduction to Bioinformatics Analysis. Curr Protoc Mol Biol 127: e92

Kalisky T, Oriel S, Bar-Lev TH, Ben-Haim N, Trink A, Wineberg Y, Kanter I, Gilad S, Pyne S (2018) A brief review of single-cell transcriptomic technologies. Brief Funct Genomics 17: 64–76

Kivioja T, Vaharautio A, Karlsson K, Bonke M, Enge M, Linnarsson S, Taipale J (2011) Counting absolute numbers of molecules using unique molecular identifiers. Nat Methods 9: 72–74

Klein AM, Mazutis L, Akartuna I, Tallapragada N, Veres A, Li V, Peshkin L, Weitz DA, Kirschner MW (2015) Droplet barcoding for single-cell transcriptomics applied to embryonic stem cells. Cell 161: 1187–1201

Lambrechts D, Wauters E, Boeckx B, Aibar S, Nittner D, Burton O, Bassez A, Decaluwe H, Pircher A, Van den Eynde K et al (2018) Phenotype molding of stromal cells in the lung tumor microenvironment. Nature medicine 24: 1277–1289

Li H, van der Leun AM, Yofe I, Lubling Y, Gelbard-Solodkin D, van Akkooi ACJ, van den Braber M, Rozeman EA, Haanen J, Blank CU et al (2019) Dysfunctional CD8 T Cells Form a Proliferative, Dynamically Regulated Compartment within Human Melanoma. Cell 176: 775–789 e718

Love MI, Huber W, Anders S (2014) Moderated estimation of fold change and dispersion for RNA-seq data with DESeq2. Genome Biol 15: 550

Luecken MD, Theis FJ (2019) Current best practices in single-cell RNA-seq analysis: a tutorial. Mol Syst Biol 15: e8746

Macosko EZ, Basu A, Satija R, Nemesh J, Shekhar K, Goldman M, Tirosh I, Bialas AR, Kamitaki N, Martersteck EM et al (2015) Highly Parallel Genome-wide Expression Profiling of Individual Cells Using Nanoliter Droplets. Cell 161: 1202–1214

Mereu E, Lafzi A, Moutinho C, Ziegenhain C, McCarthy DJ, Alvarez-Varela A, Batlle E, Sagar, Grun D, Lau JK et al (2020) Benchmarking single-cell RNA-sequencing protocols for cell atlas projects. Nat Biotechnol 38: 747–755

Natarajan KN, Miao Z, Jiang M, Huang X, Zhou H, Xie J, Wang C, Qin S, Zhao Z, Wu L et al (2019) Comparative analysis of sequencing technologies for single-cell transcriptomics. Genome Biol 20: 70

Naveed A, Cooper JA, Li R, Hubbard A, Chen J, Liu T, Wilton SD, Fletcher S, Fox AH (2021) NEAT1 polyA-modulating antisense oligonucleotides reveal opposing functions for both long non-coding RNA isoforms in neuroblastoma. Cellular and molecular life sciences : CMLS 78: 2213–2230

Osorio D, Cai JJ (2021) Systematic determination of the mitochondrial proportion in human and mice tissues for single-cell RNA-sequencing data quality control. Bioinformatics (Oxford, England) 37: 963–967

Phipson B, Zappia L, Oshlack A (2017) Gene length and detection bias in single cell RNA sequencing protocols. F1000Res 6: 595

Picelli S, Bjorklund AK, Faridani OR, Sagasser S, Winberg G, Sandberg R (2013) Smart-seq2 for sensitive full-length transcriptome profiling in single cells. Nat Methods 10: 1096–1098

Ramskold D, Luo S, Wang YC, Li R, Deng Q, Faridani OR, Daniels GA, Khrebtukova I, Loring JF, Laurent LC et al (2020) Author Correction: Full-length mRNA-Seq from single-cell levels of RNA and individual circulating tumor cells. Nat Biotechnol 38: 374

Robinson JT, Thorvaldsdottir H, Winckler W, Guttman M, Lander ES, Getz G, Mesirov JP (2011) Integrative genomics viewer. Nat Biotechnol 29: 24–26

Salcher S, Sturm G, Horvath L, Untergasser G, Kuempers C, Fotakis G, Panizzolo E, Martowicz A, Trebo M, Pall G et al (2022) High-resolution single-cell atlas reveals diversity and plasticity of tissue-resident neutrophils in non-small cell lung cancer. Cancer cell 40: 1503–1520 e1508

Schupp JC, Adams TS, Cosme C, Jr., Raredon MSB, Yuan Y, Omote N, Poli S, Chioccioli M, Rose KA, Manning EP et al (2021) Integrated Single-Cell Atlas of Endothelial Cells of the Human Lung. Circulation 144: 286–302

See P, Lum J, Chen J, Ginhoux F (2019) Corrigendum: A Single-Cell Sequencing Guide for Immunologists. Front Immunol 10: 278

Shum EY, Walczak EM, Chang C, Christina Fan H (2019) Quantitation of mRNA Transcripts and Proteins Using the BD Rhapsody Single-Cell Analysis System. Advances in experimental medicine and biology 1129: 63–79

Squair JW, Gautier M, Kathe C, Anderson MA, James ND, Hutson TH, Hudelle R, Qaiser T, Matson KJE, Barraud Q et al (2021) Confronting false discoveries in single-cell differential expression. Nature communications 12: 5692

Tang F, Barbacioru C, Wang Y, Nordman E, Lee C, Xu N, Wang X, Bodeau J, Tuch BB, Siddiqui A et al (2009) mRNA-Seq whole-transcriptome analysis of a single cell. Nat Methods 6: 377–382

Traag VA, Waltman L, van Eck NJ (2019) From Louvain to Leiden: guaranteeing well-connected communities. Sci Rep 9: 5233

Virshup I, Rybakov S, Theis FJ, Angerer P, Wolf FA (2021) anndata: Annotated data. bioRxiv: 2021.2012.2016.473007

Wang X, He Y, Zhang Q, Ren X, Zhang Z (2021) Direct Comparative Analyses of 10X Genomics Chromium and Smart-seq2. Genomics Proteomics Bioinformatics 19: 253–266

Wilusz JE, JnBaptiste CK, Lu LY, Kuhn CD, Joshua-Tor L, Sharp PA (2012) A triple helix stabilizes the 3’ ends of long noncoding RNAs that lack poly(A) tails. Genes & development 26: 2392–2407

Wolf FA, Angerer P, Theis FJ (2018) SCANPY: large-scale single-cell gene expression data analysis. Genome Biol 19: 15

Xu C, Lopez R, Mehlman E, Regier J, Jordan MI, Yosef N (2021) Probabilistic harmonization and annotation of single-cell transcriptomics data with deep generative models. Mol Syst Biol 17: e9620

Yamawaki TM, Lu DR, Ellwanger DC, Bhatt D, Manzanillo P, Arias V, Zhou H, Yoon OK, Homann O, Wang S et al (2021) Systematic comparison of high-throughput single-cell RNA-seq methods for immune cell profiling. BMC Genomics 22: 66

Zhang Q, He Y, Luo N, Patel SJ, Han Y, Gao R, Modak M, Carotta S, Haslinger C, Kind D et al (2019a) Landscape and Dynamics of Single Immune Cells in Hepatocellular Carcinoma. Cell 179: 829–845 e820

Zhang X, Li T, Liu F, Chen Y, Yao J, Li Z, Huang Y, Wang J (2019b) Comparative Analysis of Droplet-Based Ultra-High-Throughput Single-Cell RNA-Seq Systems. Molecular cell 73: 130–142 e135

Zhao Q, Wang J, Levichkin IV, Stasinopoulos S, Ryan MT, Hoogenraad NJ (2002) A mitochondrial specific stress response in mammalian cells. The EMBO journal 21: 4411–4419

Zheng C, Zheng L, Yoo JK, Guo H, Zhang Y, Guo X, Kang B, Hu R, Huang JY, Zhang Q et al (2017a) Landscape of Infiltrating T Cells in Liver Cancer Revealed by Single-Cell Sequencing. Cell 169: 1342–1356 e1316

Zheng GX, Terry JM, Belgrader P, Ryvkin P, Bent ZW, Wilson R, Ziraldo SB, Wheeler TD, McDermott GP, Zhu J et al (2017b) Massively parallel digital transcriptional profiling of single cells. Nature communications 8: 14049

Ziegenhain C, Vieth B, Parekh S, Reinius B, Guillaumet-Adkins A, Smets M, Leonhardt H, Heyn H, Hellmann I, Enard W (2017) Comparative Analysis of Single-Cell RNA Sequencing Methods. Molecular cell 65: 631–643 e634

